# Epi4Ab: A data-driven prediction model of conformational epitopes for specific antibody VH/VL families and CDR H3/L1 sequences

**DOI:** 10.1101/2025.03.13.642979

**Authors:** Nhan Dinh Tran, Krithika Subramani, Chinh Tran-To Su

## Abstract

Antibodies recognize antigens via complementary and structurally dependent mechanisms. Therefore, inclusion of antibody inputs is crucial for accurate epitope prediction. Given limited availability of antibody-antigen complex structures, it is necessary that any epitope prediction model requires minimal yet sufficient antibody inputs to ensure precise epitope identification. To address this need, we introduce Epi4Ab, an antibody-specific epitope prediction model, which focuses on identifying unique in-contact antigen residues for a given antibody. Epi4Ab requires minimal antibody inputs, specifically VH/VL families and CDR H3/L1 sequences.

## INTRODUCTION

Therapeutic antibodies are crucial in disease detection and treatment, especially for cancer and infectious diseases^1^. While effective, monoclonal antibodies used in targeted therapies are the key factor contributing to the rising cost burden over the years^2^. Antibody discovery technologies against a target have been blooming via intensive experimental and/or computational-aided efforts^3-5^. The processes are, however, laborious, resource- and time-consuming. Moreover, the designed antibodies for treatment must effectively target their antigens while also ensuring safety to succeed as therapeutics^6^. However, resistance to existing drugs is not uncommon^7^. To balance the requirements of effective therapeutics and the expenses associated with developing new drugs from scratch, drug repurposing is essential to re-adapt approved therapeutics to address mutated or newly emerging drug targets. This goal gives rise to our “reverse” approach to first identify antibody-binding region (epitope) on a new antigen given a specific antibody, initiating a pipeline of antibody engineering for the therapeutics repurposing.

An epitope is a region on an antigen defined by a linear polypeptide chain or constituted of discontinuous residual segments forming conformational patches that are complementarily bound by an antibody. Epitope prediction is a computational approach to identify the antigen regions, either linear or conformational, that could be recognized by antibodies. In contrast to generic epitope prediction, which predicts epitopes targetable by all possible antibodies and requires only antigen as input, antibody-specific epitope prediction is more targeted and necessitates both antigen and antibody as starting points.

The generic epitope prediction approach is elicited by sequence-based and structure-based methods. The former exploits advantages of extensive protein sequence data available, using deep learning-based techniques facilitated with protein language models to identify sequence patterns and predict epitopes on new antigen sequences, as demonstrated in BepiPred 3.0 ^8^, SEMA 2.0 ^9^, EpiDope^10^, and EpiBERTope^11^. On the other hand, structure-based methods rely on physicochemical characteristics inherent in the antigen-antibody interactions that was embedded in both the bound conformations of the partners. The structural features are encoded and decoded in supervised manners to make prediction on new antigen structures, as exemplified in ElliPro^12^, epitope3D^13^, SEMA 2.0 ^9^, SEPPA 3.0 ^14^, and DiscoTope 2.0 ^15^. In either scenario, accurately predicting epitopes remain challenging^16^ due to ambiguity of true epitopes as well as limited availability of the antibody-antigen complex structures.

Only a limited number of antibody-specific epitope prediction tools have been developed, requiring both antigen and antibody as inputs, such as EpiPred^17^, EpiScan^18^, and SEPPA-mAb^19^. These methods identify surface patches that could be antigenic and targeted by the given antibody. Within their own benchmarking, each method demonstrated reasonable performance and ability to distinguish epitope and non-epitope residues. Nonetheless, false positive rate remains high, leaving rooms for more developments.

An antibody binds with high affinity to its antigen via networks of weak and non-covalent interactions. The antibody-antigen interacting interface is formed by structural complementarity of the two partners’ binding regions, and this recognition is specific to the antigen.^20,21^ The antigen recognition specificity is determined by structural combinations of antibody elements such as complementarity-determining regions (CDRs) on both heavy and light chain variable domains (VH and VL), see Supplementary Figure S1. Previous studies^22-28^ have shown manipulated combinations of VH-VL frameworks (FWRs) and CDRs could allosterically modulate distinguishingly the antigen bindings, suggesting that these specific antibody structural characteristics play a substantial role in the epitope identification on the antigen.

Like EpiScan, which leverages on such heavy/light chain FWRs and CDRs information via the inputted antibody sequences for epitope prediction, we incorporate yet minimize the requirements of antibody details and concentrate on residual interaction networks embedded in the bound antigen, aiming for (i) less dependency on prior knowledge of the given antibody (which is occasionally limited) and (ii) identification of potential unique in-contact residues on the targeted antigen with the specific antibody.

To do so, we introduce Epi4Ab, a data-driven graph-based model designed to predict conformational epitopes for specific antibody VH-VL and CDRs. Epi4Ab primarily focuses on identifying antigen residues that directly interact with the given antibody, requiring only minimal antibody inputs, specifically VH/VL families and CDR H3/L1 sequences, for more efficient epitope prediction.

## METHODOLOGY

### Epi4Ab model

The Epi4Ab model learns structural interaction patterns between antigen and antibody to predict potential antibody-interacting residues (epitope) on a new antigen, using minimal input of a certain specific antibody, such as heavy-light chain variable VH-VL families and CDR H3/L1 sequences. This model is based on a multilayer Graph Neural Network (GNN) integrated with an *attention* mechanism to specifically capture distinct residual structural features embedded in the bound conformation of the antigen (Figure 1).

**Figure 1.**
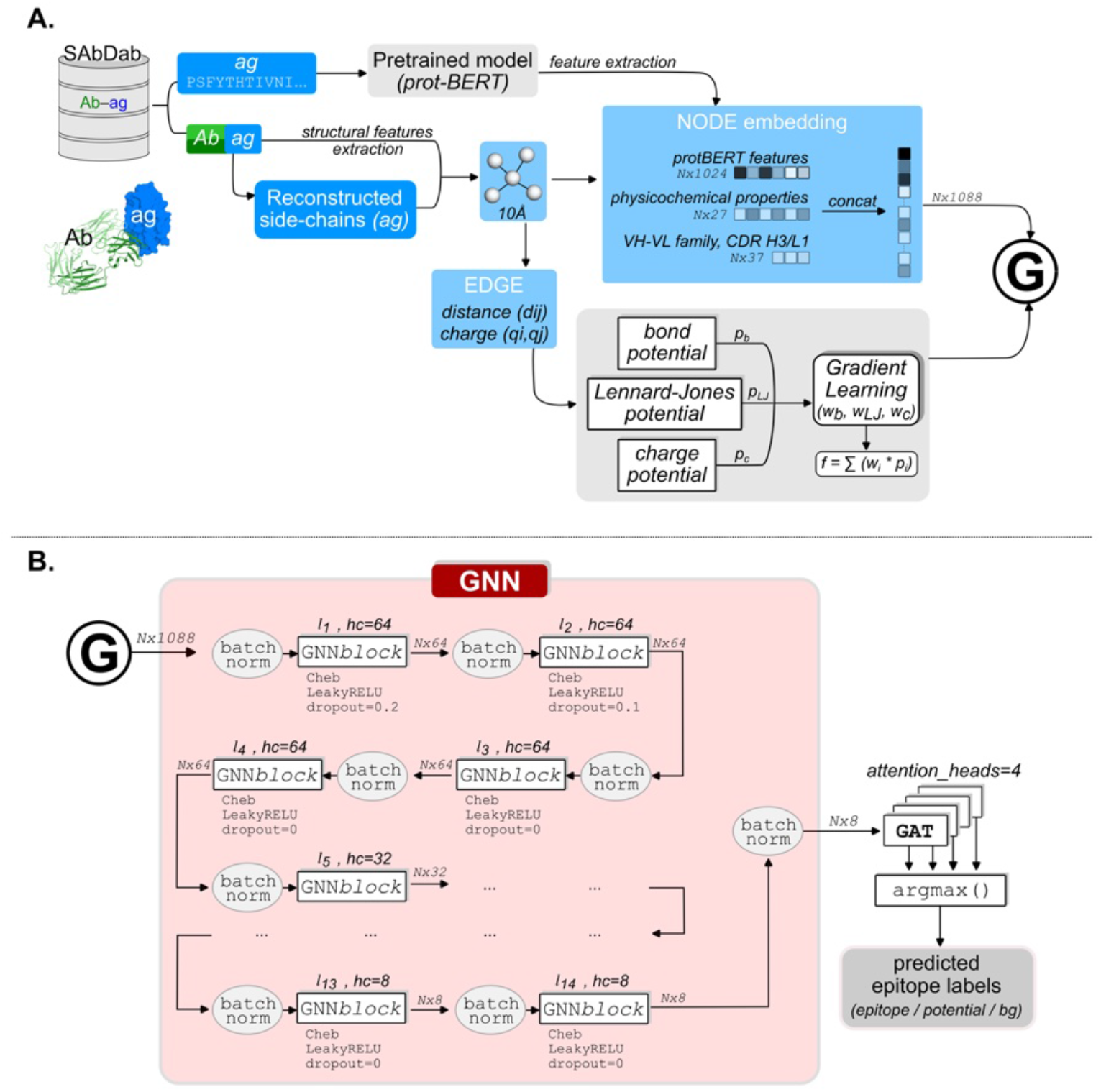
Epi4Ab workflow. (A). The model’s highlights include (i) leveraging on sequence features from the pretrained model protBERT^29^ and (ii) integrating a meta-model of learning molecular interaction potentials (bond, Lennard Jones, and charge) into the residual graph G of the antigen. (B) demonstrates the Epi4Ab’s core consisting of the Graph Neural Network (GNN) implemented with Graph Attention Network (GAT) architecture. Different numbers of layers l_i_ and hidden channels (hc) were optimized, using tensors of “number of residues N × feature size” dimension.

The Epi4Ab model leverages on a pretrained model protBERT^29^ to extract sequence patterns of antigens that interacted with their antibody partners. Simultaneously, structural features were derived using the antibody-bound conformations of the antigens. These bound antigen structures were then relaxed by reconstructing the side-chains to (i) partially mimic unbound conformation of the antigen and to (ii) augment data for subsequent model training. Antibody VH-VL families and CDR H3/L1 sequences were concurrently recorded. All the features were used for the graph construction. To enhance dynamics of residual interactions in the constructed graphs, we developed a meta-model to learn and optimize a potential scoring function, accommodating interaction potentials such as bond, charge, and Lennard-Jones potentials.

The residual graph *G* representing the antigens were fed to a multilayer GNN facilitated with Graph Attention Network^30^ (GAT) architecture to classify each antigen residue as background *non-epitope* (label 0), antibody-interacting residue or *epitope* (label 1), and *potential epitope* (label 2). To investigate effectiveness of the attention mechanism, we also implemented a naïve Graph Convolutional Network (GCN) without the GAT for comparison. Details are as below.

### Training and testing datasets

Antibody-antigen (Ab-ag) complexes were retrieved from the Structural Antibody Database^31^ (SAbDab), accessed in November 2022. Only the high resolution (≤ 4Å) Ab-ag complexes containing both antibody heavy and light chain variable domains (such as paired Fab or Fv) bound with a single chain protein antigen (length ≤ 640 residues, given our limited computational resource) were selected, resulting in 2,211 complexes. We allocated all 12 HER2 antigens from this samples into a reserved test set, and the remaining (2,199 antigens) for training sets, ensuring no HER2 antigens were included in the training or validation sets.

### Characterize structural embedded antibody-antigen interaction features

We characterized the antibody-antigen interactions by examining physicochemical properties of their binding interfaces, particularly reflected on each of the antigen residues. For each antibody-antigen complex, the bound conformation of the antigen was used to presume the structural dependency of the residual interaction network within the antigen structure (or allostery involved, if any) when bound by the corresponding antibody. The structural features included solvent exposures (using all-atom relative solvent accessibility, *rsa*, calculated by Naccess^32^ with default probe size=1.4 Å), partial charges calculated using PDB2PQR^33^, and dihedrals including phi, psi, omega, and chi angles of each residue using MDAnalysis^34^. For each antigen residue, several biophysical properties such as residue weight and volume (by IMGT^35^), hydrophobicity score (based on Kyte and Doolittle scale^36^), numbers of atoms, isoelectric point^37^ were also recorded.

Subsequently, a process of reconstructing sidechain (using FoldX5^38^) was performed for each bound conformation of the antigens to partially mimic their unbound conformations, from which the physicochemical features were re-quantified for data augmentation.

### Antibody heavy and light variable domain (VH-VL) families and CDR H3/L1 features

The VH-VL families were recorded from the retrieved SAbDab dataset, including existing families such as heavy: [IGHV01, IGHV02, IGHV03, IGHV04, IGHV05, IGHV06, IGHV07, IGHV08, IGHV09, IGHV14], and light: [IGKV01, IGKV02, IGKV03, IGKV04, IGKV05, IGKV06, IGKV08, IGKV10, IGKV12, IGKV13, IGKV14, IGLV01, IGLV02, IGLV03, IGLV06]. For those families with few entries or not known, “`others`” and “`unknown`” were recorded, respectively.

Similarly, sequences of all 6 CDR loops were extracted. For any missing CDR information, manual CDR search was performed on the SAbDab webapp using corresponding PDB entry following Chothia CDR definition. The CDR lengths were calculated accordingly.

### VH-VL-H3_len_-L1_len_ Clustering

The Ab-ag complexes were first clustered accordingly to the antibody paired VH-VL families and CDR H3/L1 lengths. Sequence global alignment using MAFFT^39^ with 1000 iterates was performed for each H3 and L1 sequence within each cluster. An in-house python script was used to estimate the alignment score using BLOSUM62 and the defaulted MAFFT *gap_score* = -1.53. The H3 and L1 sequences with the highest score were extracted as templates, representing the corresponding VH-VL-H3_len_-L1_len_ cluster. The templates were used to estimate the *H3score and L1score* features for each of the CDR H3/L1 sequences in the HER2 test set.

### Ground truth residue labels for supervised learning

We classified the antigen residues such as background non-epitope (label “0”), direct antibody-interacting residue as *epitope* (label “1”) and residues potentially targeted by other antibodies as *potential epitopes* (label “2”).

The label “1” residues were defined as the antibody-antigen interfacial contact residues within a cut-off distance of 5 Å and determined by CIPS^40^. The remaining residues were labelled as “0” (background). To lessen the resulting imbalance in the dataset labels, we introduced the label “2” as other potential epitopes.

Two generic epitope prediction tools such as Ellipro (structure-based) and BepiPred 3.0 (sequence-based) were employed to first identify potential antigenic surface residues. The two methods were selected due to their largest coverage among several available methods^16^ to filter out background non-epitope residues. Consensus of the resulting antigenic surface residues (excluding those initiated with label “1”) by the two methods were then defined as label “2” residues.

### Residual Graph construction

Each antigen was represented by a residual graph *G* = {*V, E*} with nodes *V* representing the antigen residues and edges *E* representing interactions between the residues. A cut-off Cα-Cα distance of 10 Å was used to determine neighboring nodes for each node *v*_*i*_. Each node *v*_*i*_ was represented by a concatenated tensor including the characterized sequence and structural features together with the respective antibody records of the VH-VL families and CDRs (Figure 1).

To enhance dynamics of the graph, we integrated several potentials estimating the edge attributes and hence mimicking the biophysical interactions within the networks. These include bond (*p*_*b*_), Lennard-Jones (*p*_*LJ*_), and charge (*p*_*c*_) potentials incorporated in a scoring function as below:

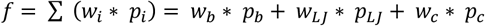

In which *w*_*b*_, *w*_*LJ*_, and *w*_*c*_ are the respective weighted coefficients and were optimized via an implemented gradient learning meta-model (Figure 1). This process was supervised and performed using 5-fold cross validations on the full training sets. The set of weights that resulted in the highest F1_epitope_ score in the validation sets and least over-fitting was selected for further training.

The three potentials were defined as below:

(i) Bond mimicking potential: 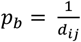
(ii) Lennard-Jones mimicking potential: 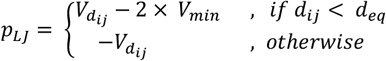 With 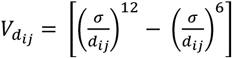 representing potential energy at distance_*ij*_ between two residues *i,j* (Cα-Cα) and *σ* = 3.4 Å is the distance at which 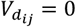. V_*min*_ *≈ ™*0.25, representing potential energy at the equilibrium 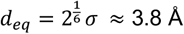.
(iii) Charge potential: 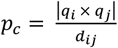, in which *q*_*i*_, *q*_*j*_ are partial charges of residues *i, j*

### The GNN architecture

The Graph Neural Network (GNN) was implemented with 14 layers, each of which employed a Chebyshev spectral graph convolutional operator^41^ (Cheb) and activated using LeakyRELU. Feature dropouts (ratio of 20% and 10%) were used for the first two layers. Different numbers of hidden channels were used subsequently for the layers, i.e. 64×4→32×4→16×4→8×2. Other parameters such as filter_size=5, batch_size=50 and “BatchNorm” were used. The model was trained for 500 epoches with learning_rate = 0.0005, using Adam optimizer and with num_attention_head=4.

Probability (*prob*) was estimated for each label = [0,1,2], the largest of which determined the predicted label outcome for each antigen residue. The predicted probability was then converted into prediction score as following: 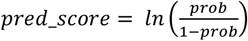.

### Benchmarking against several epitope prediction methods

The test set of 12 HER2 antigens was used for epitope prediction on several webservers with default settings such as epitope3D^13^, SEMA 2.0^9^, DiscoTope 3.0^42^, SEPPA 3.0^14^, and SEPPA-mAb^19^. Given no available webserver and due to incompatibility in the published EpiScan’s source code, 8 non-redundant tested antigens provided by EpiScan^18^ (which excluded from our training set) was used to test against our Epi4Ab.

To maintain competitiveness across all the methods, we re-evaluated true positives using a predefined cavity sphere (green circle in Figure 2) that encompasses the antibody-antigen interface in the ground truth. Therefore, a predicted epitope residue (by any of the tools) was considered as a true positive only if it was located within this sphere and directly interacted with the antibody, defined by CIPS^40^. Recall, Precision, and F1_epitope_ score were used as metrics to compare across the methods.

**Figure 2.**
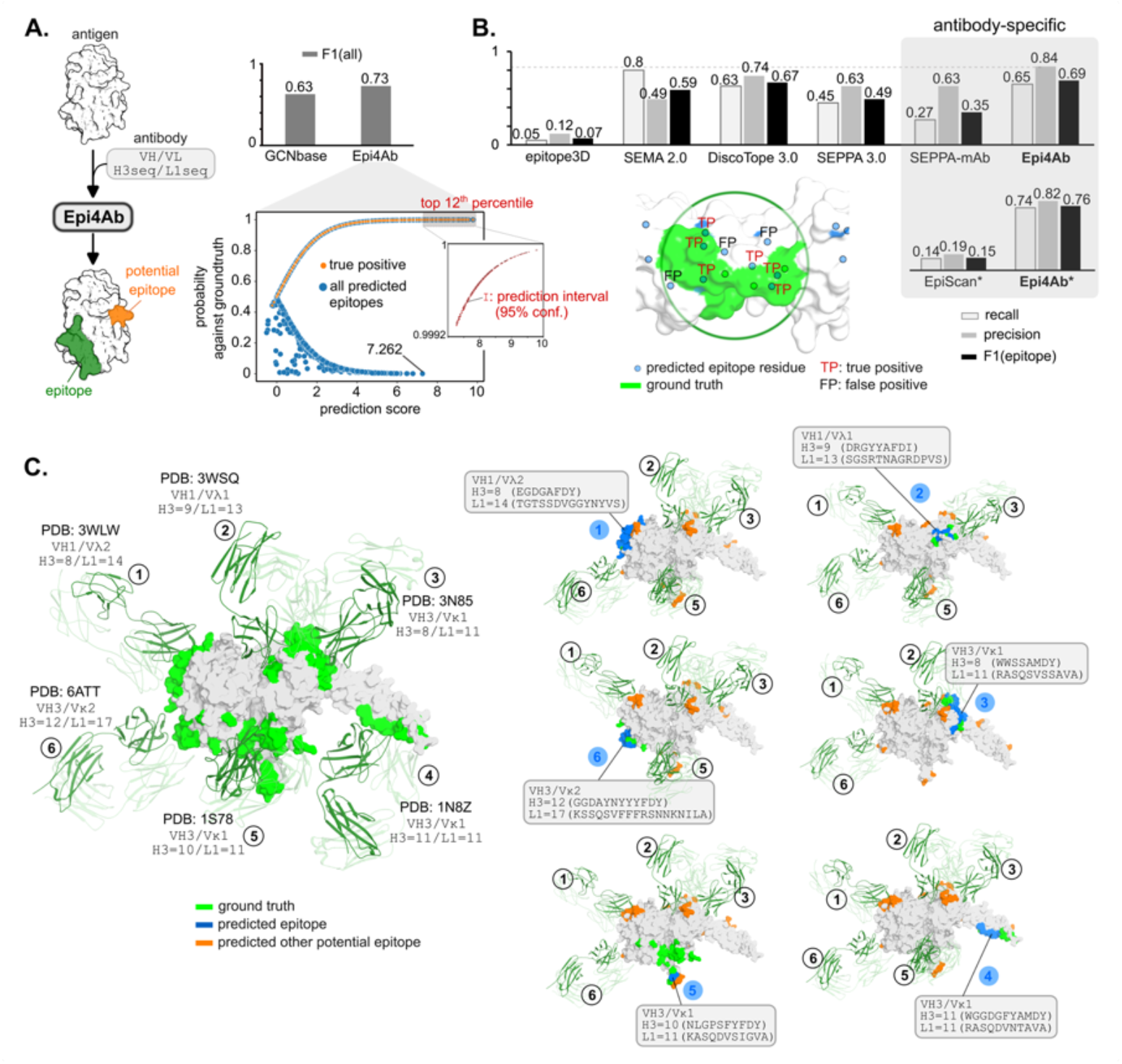
Performance of Epi4Ab. (A). Overview of the Epi4Ab model. The “attention” mechanism enhanced the epitope identification, with highest prediction scores (>7.262) ranked in the top 12^th^ percentile with 95% confidence. (B). The Epi4Ab model outperforms across various generic and antibody-specific epitope prediction methods. When compared with EpiScan*, the test set containing 8 non-redundant antigens suggested by EpiScan^18^ was used for compatibility. For benchmarking consistency, true/false positives were re-evaluated (see Methods), by implementing a (green) cavity sphere encompassing the true antibody-antigen interaction interface. (C). Structural presentations of the Epi4Ab predictions using the HER2 test set. To simplify, only those HER2-binding antibodies with the least epitope overlaps are represented: ground truth (green), predicted epitopes (blue) and potential epitopes (orange). HER2 is shown in gray surface and the antibodies in dark/light green cartoons.

## RESULTS

### Epi4Ab predicts conformational epitopes given minimal antibody input

To address limitations and occasional unavailability of antibody structures, the Epi4Ab model was trained additionally with minimal antibody input, specifically VH-VL families and CDR H3/L1 sequences, to identify antibody-specific epitopes on antigens. We defined “epitope” as direct antibody-interacting residues on the antigen targeted by the specific antibody, distinguishing the predicted *epitopes* from the *potential epitopes* (regions potentially targeted by other antibodies) in our results.

Since interaction networks of epitope residues differed from those of neighboring non-epitope residues, an *attention* mechanism was incorporated into our graph neural network (GNN) model to focus on the distinct structural embedded features. Compared to our baseline model (a naïve GCN without the attention), adding the *attention* improved the prediction accuracy and increased true positives (reflected in F1 score) estimated on the 12 HER2 antigens in the test set (Figure 2A). Repeated testing results (on 10-fold cross validation) affirmed the highest prediction score (threshold >7.262) ranked in the top 12^th^ percentile, with 95% confidence.

We used this HER2 test set to benchmark our Epi4Ab model against various available generic epitope prediction tools (webservers) such as epitope3D^13^, SEMA2.0^9^, DiscoTope3.0^42^, and SEPPA3.0^14^ and two recent antibody-specific epitope prediction methods SEPPA-mAb^19^ and EpiScan^18^. To maintain competitiveness and avoid biases among the tools, we re-evaluated the *true positives* for each tool’s predictions, i.e. only the predicted epitope residues that were located within the cavity sphere (green circle in Figure 2B) encompassing the antibody-interacting residues and that directly interacted with the corresponding antibody. Results across the tools indicated that Epi4Ab outperformed (ROC_AUC=0.81 and PR_AUC=0.59, shown in Table 1), achieving the highest precision (0.84) and F1_epitope_ (0.69). Among these generic epitope prediction tools, DiscoTope 3.0 performed comparably well, ranking the second best (precision=0.74, F1_epitope_=0.67) in the comparison list (Figure 2B).

**Table 1:** Epi4Ab benchmarking against various generic and antibody-specific epitope prediction tools, estimating on ROC_AUC (area under the ROC curve), PR_AUC (area under Precision-Recall curve), and Accuracy.

| Epitope prediction model |  | ROC_AUC | PR_AUC | Accuracy |
| --- | --- | --- | --- | --- |
| Generic | epitope3D | $0.43 \pm 0.03$ | $0.37 \pm 0.06$ | $0.53 \pm 0.06$ |
| | SEMA 2.0 | $0.63 \pm 0.07$ | $0.45 \pm 0.07$ | $0.60 \pm 0.07$ |
| | DiscoTope 3.0 | $0.75 \pm 0.10$ | $0.61 \pm 0.15$ | $0.79 \pm 0.07$ |
| | SEPPA 3.0 | $0.62 \pm 0.06$ | $0.47 \pm 0.06$ | $0.67 \pm 0.07$ |
| Antibody-specific | SEPPA-mAb | $0.59 \pm 0.09$ | $0.47 \pm 0.11$ | $0.68 \pm 0.09$ |
|  | **Epi4Ab** | **$0.81 \pm 0.09$** | **$0.59 \pm 0.17$** | **$0.91 \pm 0.03$** |
| | EpiScan* | $0.49 \pm 0.03$ | $0.47 \pm 0.09$ | $0.50 \pm 0.09$ |
|  | **Epi4Ab\*** | **$0.85 \pm 0.07$** | **$0.66 \pm 0.13$** | **$0.91 \pm 0.04$** |

\*Epi4Ab was tested against 8 non-redundant antigens in the test set used by EpiScan

When comparing with the antibody-specific tool EpiScan, we instead used the non-redundant test set (8 antibody-antigens complexes) suggested by EpiScan to test against our Epi4Ab. These 8 antigens were excluded from our training set, hence being fully “unseen” to our Epi4Ab model. When testing on the ability to identify direct antibody-interacting residue, our Epi4Ab performed better (F1_epitope_=0.76 vs 0.15). Similarly, Epi4Ab could also identify the interacting residues better than SEPPA-mAb on the HER2 test set (F1_epitope_=0.69 vs 0.35). This suggests an advantage in identifying direct interacting residues over surface patches resulted by the two recent methods.

The HER2 antigen is recognized at different binding interfaces by multiple antibodies comprising various combinations of VH-VL families and CDRs sequences (Figure 2C, left). Given the respective antibody input, the Epi4Ab accurately identified the corresponding interfacial patch with precise interacting residues, and in fact, effectively distinguished the non-epitope residues (average precision approximately 0.97). Interestingly, the resulting predicted *potential epitope* regions (orange in Figure 2C) were found overlapping with the binding interfaces by other antibodies, indicating the interpretation ability of the model given the embedded structural features within the bound conformation of the antigen.

### Does the minimal antibody input impact the Epi4Ab epitope prediction or just merely add supplementary features?

Previous studies^25,27,28,43^ demonstrated antibody heavy/light chain variable FWRs and CDRs modulated antigen binding through allosteric effects. This elucidated the associated role of the variable (VH/VL) family, as determined by FWRs, and CDRs in identifying the interaction interface on antigens; hence, these antibody features are necessary to predict interfacial residues on the antigen using the Epi4Ab approach.

The 12 antibodies that bound to the tested HER2 antigens contain varying VH-VL family combinations such as VH1/VH3 (heavy) paired with Vκ1/Vκ2/Vλ1/Vλ2 (light) and different CDRs (particularly with diverse H3 and L1 lengths) and interacted with the HER2 at different epitopes (Figure 2C and 3A). To assess the model sensitivity to the antibody input, we performed the Epi4Ab inference on the HER2 structures individually using 472 unique VH-VL-H3_len_-L1_len_ combinations available in the dataset. To further increase the variances among the simulated inputs, the most differentiated H3 and L1 sequences from the cluster consensus were selected with respect to each of these VH-VL-H3_len_-L1_len_ combinations. The hypothesis stated that if the antibody input was irrelevant or having minute effect on the epitope prediction of the Epi4Ab, the changes in the model’s ability to identify the antibody-interacting residues would remain insignificant. We used F1_epitope_ to reflect this ability, emphasizing on identifying the antibody-interacting residues (true positives) on each HER2 antigen.

The results indicated that the calculated F1_epitope_ scores varied noticeably given the VH-VL-H3-L1 input changes for each of the tested HER2 antigen structure, with the most substantial variation observed in the cases of 3N85 (Figure 3B, left). The antibody-binding cavity in 3N85 exhibited similar physicochemical properties to those found in other test cases such as 3H3B and 6BGT (Figure 3B, right), e.g. slightly exposed (mean relative solvent accessibility ∼19-23%) and large (volume >2800 Å^3^); however, the epitope identification within this cavity surprisingly showed a broader deviation due to the various VH-VL-H3-L1 inputs; in fact, abrogated in some instances. Noticeable changes were also observed in other test cases, the antibody of which contain paired VH3-Vκ1 domains such as 1N8Z, 3BE1, 6BGT, and 7MN8. This suggests the antigen regions recognized by these common VH3-Vκ1 pairs would be more susceptible to antibody manipulation.

**Figure 3.**
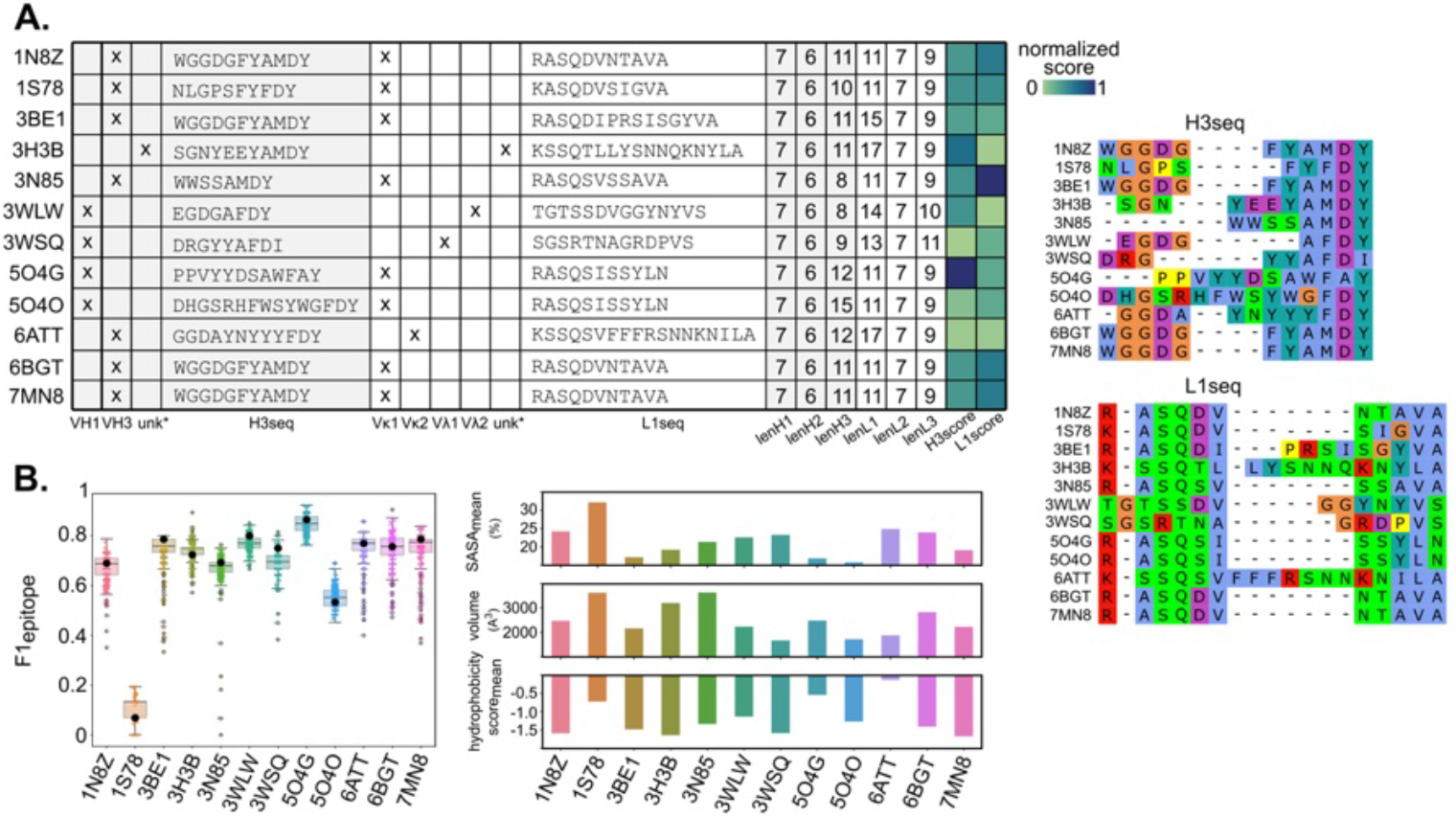
Effect of the VH-VL-H3-L1 input changes on the epitope prediction of the Epi4Ab model. (A). The VH-VL and CDRs input of the 12 antibodies that target the HER2 antigens in the test set. The H3 and L1 sequences were aligned to the respective consensus template, the VH-VL-H3_len_-L1_len_ cluster of which the antibody was classified. The alignment scores were normalized, with the highest (=1) indicating the most similarity to the template. Multiple sequence alignment of the 12 sequences of CDR H3 and L1 were performed using MAFFT^39^ to highlight their differences. The residues are colored using Clustal scheme. (B). The epitope inference results of each tested HER2 structure given various VH-VL-H3-L1 inputs. The starting F1_epitope_ scores corresponding to the original VH-VL-H3-L1 combination in each tested HER2 are shown in black dots. The antibody-binding cavities in each of the original tested HER2 are demonstrated using several main physicochemical features such as solvent exposure (mean relative solvent accessibility), volume, and hydrophobicity score (the higher, the more hydrophobic).

Observed in the test cases involving paired VH1-Vλ* antibody, such as 3WLW and 3WSQ, the calculated F1_epitope_ scores of the latter decreased more when challenged with various input VH-VL-H3-L1 combinations. In this 3WSQ instance, the antibody light chain variable interacting with HER2 belongs to the Vλ1 family, featuring distinct CDR H3 and L1 sequences compared to the other test cases. Notably, both the CDR H3 and L1 contain additional positively charged arginine (Figure 3A). The low normalized alignment CDR H3 and L1 scores also indicated the major sequence variation against the consensus H3 and L1 sequences, respectively within the VH1-Vλ1-H3_len=9_-L1_len=13_ combination. However, these observations were not found in the cases of 3WLW with the paired VH1-Vλ2 families.

To note, there was no correlation (data not shown) observed in the retrieved dataset between the antibody CDR H3/L1 length/sequence and the other physicochemical features such as volume, solvent exposure, or hydrophobicity of the antibody-binding region on the antigens, indicating the independency of these features. Therefore, it elicited that the VH-VL-H3-L1 input was more than essential descriptors but also manipulating the epitope prediction outcomes. Since the necessary antibody inputs are minimal, it highlights the effectiveness of our Epi4Ab model in predicting the antibody-interacting residues on the given antigen.

## DISCUSSION

We developed the Epi4Ab model to predict potential antibody-interacting residue on a new antigen given certain specific antibody VH-VL families and CDR H3/L1 sequences. Reasonable performance of Epi4Ab suggests feasibility for the epitope prediction to leverage on minimal dependency on the antibody structural input, which otherwise is limited or occasionally unavailable.

We prioritized on identifying unique in-contact residues rather than conventional antigenic surface patches to enhance manipulation of antigen targeting. This would consequently downplay the secondary role of non-interacting residues located at the antibody-antigen interface. In exchange, we used bound conformations of the antigens to preserve both short- and long-ranged effects and hence maintain local interaction networks within the antigen structures when bound by antibodies. This, in fact, facilitated the Epi4Ab prediction of *potential epitope* (orange in Figure 2C), which depicted regions potentially recognized by any other antibodies. For instance, while accurately identifying epitopes on HER2 targeted by Trastuzumab (PDB: 1N8Z), Epi4Ab also predicted several residues potentially recognized by Pertuzumab (PDB: 1S78) when provided with Trastuzumab’s inputs. The two antibodies’ colocalization and co-binding to HER2 at distinguished epitopes has been studied previously^44-46^, demonstrating the possible allosteric mechanism within the HER2 structure when bound by either antibody.

Antibody and antigen binding occurs in a lock-and-key, induced fit, or involving conformational selection mechanism. In the lock-and-key model, antibody acts as the lock and attaches to the antigen (the key), without significantly altering the structural conformations of either partner. Conversely, both antibody and antigen undergo substantial changes in the induced fit model, particularly at their interaction interface, whereas the conformational selection model involves the antigen sampling certain unique conformational states prior to binding with antibody. Overall, structural dependency plays a crucial role in the antibody-antigen binding mechanism, suggesting that preserving a certain degree of the structural dependency during the model’s supervised learning process might be essential to capture the underlying conformational changes, thereby facilitating the identification of the near-native epitopes.

To accommodate the structural dependencies involving antibody interactions, current antibody-specific epitope prediction methods incorporate more comprehensive antibody inputs; for example, SEPPA-mAb requires additional bound conformation of the antibody while EpiScan employs antibody sequences. As illustrated in the test cases of EpiScan (Figure 2B), incorporating antibody sequences alone might be insufficient for accurate identification of in-contact epitope residues. Nevertheless, obtaining an accurate antibody structure, even those based on homology models, remains challenging.

Along with this context, to incorporate while minimizing the structural dependencies, Epi4Ab employed the sidechain reconstructions to “relax” the bound conformation of the antigens, partially mimicking the unbound conformation; however, the Epi4Ab model still experienced limitations due to the structural dependency. For instance, manipulating exposure of several interfacial residues or disabling fully the structural dependency (e.g. remodeling unbound antigen conformation using AlphaFold) compromised the Epi4Ab’s performance (data not shown). To address this challenge for future work, we are developing metamodels to first enhance detection of potential binding regions on the unbound antigen given the specific antibody CDR H3. The ensembles of these models would facilitate in estimating the structural dependency needed to accommodate the CDR H3 interactions, leading to more efficiently identifying the in-contact residues at the interface. This will further include the VH-VL orientations in our future developments.

Overall, our epitope prediction model for specific antibody VH-VL families and CDR H3/L1 sequences, Epi4Ab, offers a more holistic view of antibody-antigen complexes. Improved models could further clarify the role of diverse antibody VH-VL and CDR (particularly H3 and L1) combinations in antigen targeting. The identified epitopes could aid in designing antibody CDR-H3 patterns for better antigen binding, addressing a key challenge in the field.

## DATA AVAILABILITY

The source codes of Epi4Ab can be found here https://github.com/AMPMgroup/Epi4Ab.

## ACKNOWLEDGMENT

This work was supported by the National Medical Research Council grant NMRC-OFYIRG (MOH-OFYIRG20nov-0018). We thank Khin Bhone Pyae in assisting the model benchmarking. The authors used Copilot, a built-in in the Microsoft 365 (Word) to assist in language improvement.

