## Supplemental Figure S1 for "Epi4Ab: A data-driven prediction model of conformational epitopes for specific antibody VH/VL families and CDR H3/L1 sequences"

**Supplementary Figure S1**

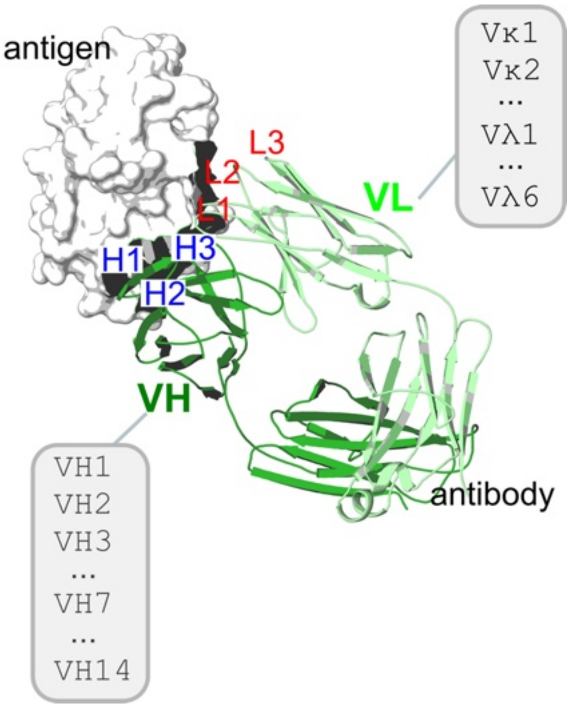


*Figure S1: Illustration of antibody-antigen complex structure, in which the interacting interface demonstrates the complementary binding between the antigen and the antibody CDR loops (heavy: H1, H2, H3, and light: L1, L2, L3). The antibody heavy/light variable domains (VH/VL) are classified into various families. Several VH/VL families recorded by the SAbDab are listed and used in the Epi4Ab's model training.*
